# Mapping densely packed GPIIb/IIIa receptors in murine blood platelets with expansion microscopy

**DOI:** 10.1101/2021.02.16.431449

**Authors:** Hannah S. Heil, Max Aigner, Sophia Maier, Prateek Gupta, Luise M.C. Evers, Vanessa Göb, Charly Kusch, Mara Meub, Bernhard Nieswandt, David Stegner, Katrin G. Heinze

**Author notes:** corresponding authors: Katrin G. Heinze, Rudolf Virchow Center for Integrative and Translational Bioimaging, University of Würzburg, Würzburg, Germany or David Stegner, Institute of Experimental Biomedicine, University Hospital Würzburg, Würzburg, Germany.

## Abstract

Interrogating small platelets and their densely packed, highly abundant receptor landscape is key to understand platelet clotting, a process that can save lives when stopping blood loss after an injury, but also kill when causing heart attack, stroke or pulmonary embolism. The underlying key receptor distributions and interactions, in particular the relevance of integrin clustering, are not fully understood is because of highly abundant and densely distributed GPIIb/IIIa receptors. This makes receptor distributions difficult to assess even by super-resolution fluorescence microscopy. Here, we combine dual-color expansion and confocal microscopy with colocalization analysis to assess platelet receptor organization without the need of a super-resolution microscope. We show that 4x expansion is highly straight-forward for super-resolution microscopy of platelets, while 10x expansion provides higher precision at the price of increased efforts in sample preparation and imaging. Quantifying various receptor colocalization scenarios we demonstrate that expansion microscopy can pinpoint receptor distributions and interactions in resting and activated platelets being superior to conventional methods that fail in such dense 3D scenarios with highly abundant receptors. We reveal the presence of GPIIb/IIIa clusters in resting platelets, which are not affected by platelet activation indicating that they contribute to the rapid platelet response during platelet clotting.

**Graphical Abstract:** 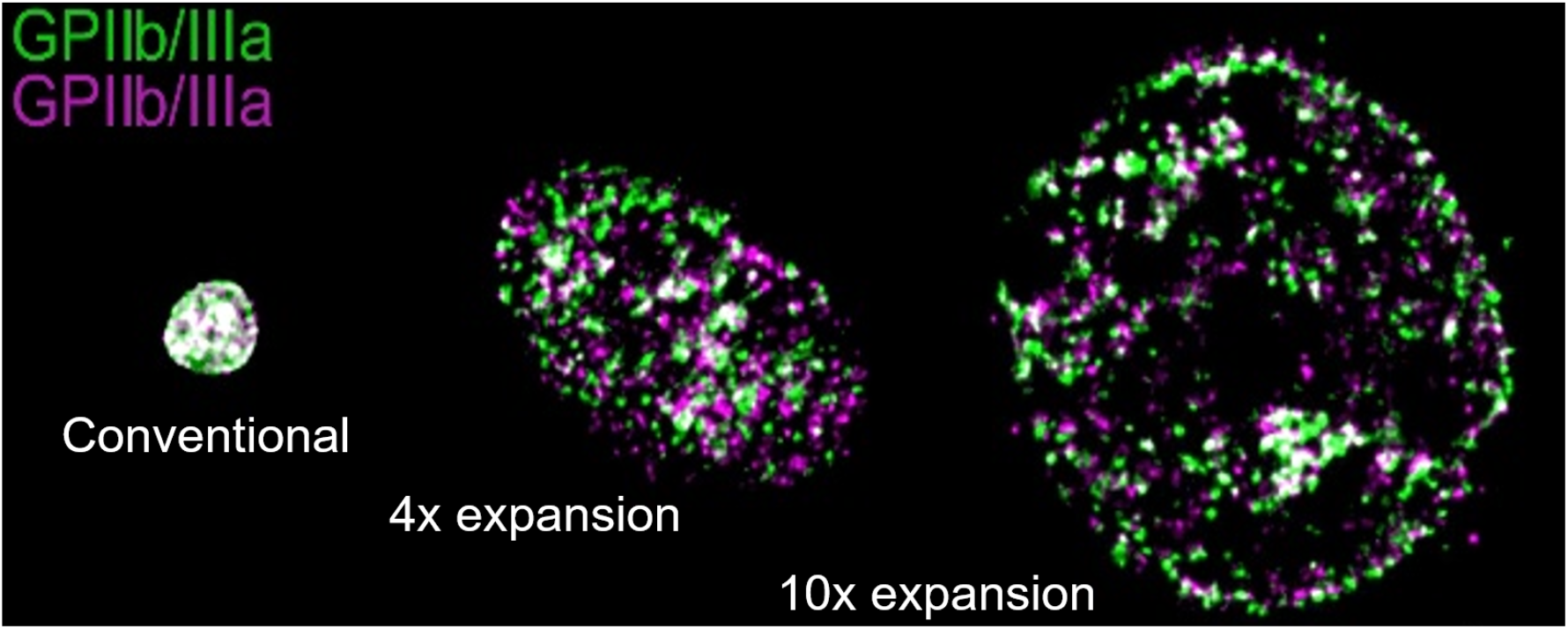

## Introduction

Integrins are a superfamily of heterodimeric cell-surface receptors that comprise an α and a β subunit.^1,2^ Each subunit consist of a large N-terminal extracellular domain that contributes to ligand binding, a single-pass transmembrane domain and a cytoplasmic tail providing binding sites for adaptors, and signaling proteins that contribute to the bidirectional integrin signaling.^3,4^ Integrin receptors are widely expressed in mammalian tissues and serve as adhesive units, which are, however, highly dynamic regulators of cellular responses. This is in particular true for platelets that require an acute and dynamic regulation of integrin affinity and ligand binding to enable rapid platelet adhesion at sites of vascular injury on the one hand and to prevent uncontrolled thrombosis of the other hand. The αIIbβ3 integrin (also named GPIIb/IIIa or CD41/CD61) is exclusively expressed by megakaryocytes and platelets and binds a number of ligands, including fibrinogen, fibrin, fibronectin and von Willebrand factor.^5^ This receptor is of high clinical relevance as GPIIb/IIIa inhibitors are widely used to prevent thrombosis and treat myocardial infarction.^6^ Moreover, GPIIb/IIIa serves as model receptor to study integrin biology.^4^ In the last decades key components of both inside-out and outside-in signaling have been identified; the first represents the functional up-regulation of the receptor’s affinity, the latter generates integrin-mediated signaling responses following ligand binding.^3,4^ In addition, it is widely accepted that clustering of integrin receptors in hetero-oligomers modulates avidity and supports the assembly of the outside-in signalosome;^7,8^ unfortunately, our current understanding of the underlying processes and the receptor distribution is far from complete.^4,9^ One reason is the high receptor density on the platelet surface. Platelets are small in size (2-3 µm in diameter) and express 50,000-100,000 copies of GPIIb/IIIa on their surface, while even more can be mobilized from intracellular stores.^10,11^ Thus, visualizing and quantifying GPIIb/IIIa distribution on the platelet surface is very challenging. Strictly spoken, a resolution in the nanometer range is required to gain an image based receptor map on the single molecule level.^12^ Electron microscopy would provide such nanometer resolution, however, despite its unrivaled resolution power, it lacks efficient staining for specific receptors and immunogold labeling is hard to achieve at saturating conditions. In contrast, fluorescence labeling is efficient and selective and thus allows visualization of single membrane receptors when combined with super-resolution imaging.^13–15^ A variety of fluorescence super-resolution modalities have been reported over the past century, each with its own advantages and limitations.^16^ Multi-color 3D single molecule localization microscopy allows to effectively decipher receptor interaction and organization at a molecular level; however, it requires a high level of expertise and equipment and fails at high molecular density scenarios^17^ such as highly abundant and dense platelet receptor distributions we are tackling here. A particularly promising, rather young method is expansion microscopy (ExM).^18^ In contrast to most other super-resolution techniques ExM does not require a high-end super-resolution microscope but uses physical volumetric expansion of the sample. Linking the fluorescently labeled proteins to a swell-able hydrogel eventually turns any conventional microscope into a super-resolution imaging device. The major downside is that isotropic expansion requires a highly processed sample and harsh preparation steps: After fixation and immunolabeling the sample structure itself needs to be tightly linked into the hydrogel network and has to be disrupted by breaking protein bonds and crosslinks through either protein digestion or degradation^19^. Depending on the hydrogel composition the gel will swell up to 10x of its original size expanding the sample to up to 1000x of its original volume^20^ (see also supplemental Figure I). With physical expansion, the labeling density decreases respectively, and the sample additionally undergoes harsh chemical treatment during gelation and digestion, which typically results in a 50% loss of fluorescent markers.^19^ Nevertheless, the resolution gain by ExM allows quantitative image-based comparison of two (fluorescently labeled) molecular distributions. Eventually, a colocalization coefficient can be calculated providing an intensity correlation measure of both color channels.^21^ The commonly used Pearson’s colocalization coefficient excels in robustness under varying signal-to-noise conditions, however fails when number of events detected in each channel are not comparable. The two Manders’ colocalization coefficients are less robust to noise, but make analysis more versatile as the approach allows to directly retrieve the overlapping fraction of one protein to the other and vice versa.^22^ In ExM reporting the overlapping fraction regardless of the degree of asymmetric signal loss in both channels is crucial.

Here we show how ExM and image analysis can be tuned to advance studies of dense and highly abundant platelet receptor interactions, which are difficult to tackle otherwise. Our quantitative approach involves a combination of ExM, confocal microscopy and colocalization analysis with optimized sample preparation for 4x and 10x expansion as well as tailored image analysis pipelines. Supported by a simulation tool for platelet receptor distributions we find that only 10x expansion provides the required resolution to distinguish colocalized from independent receptor distributions for highly abundant GPIIb/IIIa and GPIX receptors on resting and activated platelets.

## Methods

The authors declare that materials and data are available upon reasonable request from the authors. A selection of typical images and the code for simulation of platelet receptor distributions are available on ZENODO (doi: 10.5281/zenodo.4117555), where we also provide a video protocol of the sample preparation for 10x ExM. An extended version of Materials and Methods as well as detailed supplementary protocols for 4x and 10x platelet ExM are available in the Supplemental Material. All methods were carried out in accordance with relevant guidelines and regulations.

### Platelet Isolation

C57Bl/6J mice served as blood donors and animal experiments were approved by the district government of Lower Franconia (Bezirksregierung Unterfranken). Washed platelets were prepared as described elsewhere^23^. Round coverslips (Ø12 mm, #1.5, #1.5 Menzel and Ø24 mm, #1.5 Plano GmbH) were cleaned by 1 h sonication in chloroform (472476, Sigma), followed by 1 h sonication in NaOH (6771, Roth), ddH_2_O washed, dried at 70°C and stored in Ethanol (34852, Sigma). The clean coverslips were coated with 2 M glycine (A1067, AppliChem) in PBS for 10 min and washed 2x with PBS and 1x with Tyrodes buffer. The platelet suspension was deposited onto the coated coverslips (30*10^6^/ 15*10^6^ platelets for 24 mm/12 mm) and incubated for 30 min at 37°C for settling. Platelet activation was achieved by complementing the Tyrodes buffer with 2 mM Ca^2+^ and the addition of 0.01 U/ml Thrombin (10602400001, Roche) to the platelet suspension.

### Immunostaining

#### Fixation

Platelets were fixed with glyoxal solution (19.89% (wt/wt) pure ethanol, 3.15% (wt/wt) glyoxal (128465, Sigma), 0.7% (wt/wt) acetic acid (7332.1, Roth) in ddH_2_O) at pH 5.0 for 20 min ^24^ at RT, washed 5x with PBS and blocked with 5% bovine serum albumin (BSA, A3983, Sigma) in PBS for 2 h, then washed with PBS.

#### Immunolabeling

Platelets were immersed in 10 µg/ml fluorescently labeled antibody (produced in-house: IgG MWReg30 against GPIIb/IIIa, IgG JON6 against GPIIb/IIIa on different epitope, and IgG p0p6 against GPIX as described previously^25^) conjugated with

Alexa Fluor 488 (A20181, Thermo Fisher) or Alexa Fluor 594 (A20185, Thermo Fisher) solved in PBS with 5% BSA for 30 min at 37°C, ensuring saturated labeling of all available epitopes (tested with flow cytometry). Afterwards, coverslips were washed 5x with PBS. The antibodies’ degree of labelling was measured by spectrometric analysis (NanoDrop One, Thermo Fisher Scientific).

### Platelet Expansion

#### 4x expansion

(adapted from Chozinski et al 2016 ^26^). Immunolabeled platelets on coverslips were incubated at RT with linking solution (0.1 mg/ml Acryloyl-X in PBS, A20770, Invitrogen) for 12 h. A 50 µl drop of monomer solution (2 M NaCl, 2.5% (wt/wt) acrylamide (AA, A9099, Sigma), 0.15% (wt/wt) N,N′-methylenebisacrylamide (M7279, Sigma), 8.625% (wt/wt) sodium acrylate (SA, 408220, Sigma) in PBS), containing 0.2% freshly added ammonium persulfate (APS, 9592.3, Roth) and tetramethylethylenediamine (TEMED, 2367.3, Roth) was placed on elastic film (parafilm, Sigma) and covered by the sample glass coverslip (2x washed with PBS). After gelation for 2 h at RT in a humidified chamber the samples were transferred into digestion buffer (8 U/ml Proteinase K (P4850, Sigma), 50 mM Tris pH 8.0, 1 mM EDTA (ED2P, Sigma), 0.5% Triton X-100 (28314, Thermo Fisher) and 0.8 M guanidine HCl (50933, Sigma) in ddH_2_O), and incubated overnight (8 h) in a humidified chamber. After removal of the coverslip the gel was expanded in ddH_2_O for 24 hours, immobilized on a poly-D-lysine (02102694, MP Biomedicals) coated Ø24 mm, #1.5 coverslip and imaged.

*10x expansion* (adapted from Truckenbrodt *et al*. 2018 ^27^ and modified). The 10x monomer solution consists of 2.67% (wt/wt) N,N-dimethylacrylamide (DMAA, 274135, Sigma), and 0.64% (wt/wt) sodium acrylate (SA) in ddH_2_O and was bubbled with N_2_ to purge out molecular oxygen for 40 min on ice before adding 0.0036 g/ml of KPS (206224, Sigma). After another 15 min of O_2_ purging 0.4% TEMED were added, and 50 µl of final monomer solution immediately placed on elastic film and covered with the sample (on coverslip). After gelation in a N_2_-filled humidified chamber for 48 h at 4°C, digestion was performed for 12 h at 50°C.

### Fluorescence Imaging

Dual color confocal microscopy was used (Leica TCS SP5 II; 63x 1.2 NA water immersion objective, 8000 Hz resonant scanning; pixels: 1024×1024, voxels: x,y = 41 nm & z=170 nm, pinhole: 1 Airy unit, scanning: 8x line averaging, 3 frames accumulated, exc.: 488 nm & 561 nm, detection: 500-550 and 600-650 nm in sequential mode, Hybrid detector (HyD). Chromatic aberrations were corrected by distortion matrices using bUnwarpJ^28^ (calibration by 100 nm 4-colour fluorescent microspheres (TetraSpeck, T7279) as described elsewhere^29^). Images were deconvolved (Huygens Professional, SVI, Leiden, Netherlands) with a theoretical PSF and a signal-to-noise threshold of 30 counts.

### Data Analysis

Image preprocessing, colocalization analysis and statistical validation, as well as the simulation, dSTORM imaging and analysis are described in detail in the *Online Supplementary Methods*.

## Results

### Optimizing ExM for receptor colocalization on platelets

The accuracy of a colocalization study mainly depends on the spatial resolution of the underlying image.^12^ In order to compare the performance of the colocalization analysis for different resolution levels, we established two different kinds of expansion protocols leading to 4x and 10x expansion along previously published work.^19,27^ Before immunolabeling and expansion, purified resting platelets are settled on a glycine coated surface and chemically fixed without any signal compromise (supplemental Figure II A,B) to avoid unintended receptor reorganization by platelet activation. Here, glyoxal fixation outperformed PFA fixation in both, labeling efficiency and minimal background signal^24^ (supplemental Figure II C,D).

Figure 1A shows the respective deconvolved confocal images of GPIIb/IIIa receptors on unexpanded versus 4x and 10x expanded platelets. The GPIIb/IIIa receptors are targeted by monoclonal antibodies carrying *either* an Alexa Fluor 594 (Figure 1, magenta) *or* an Alexa Fluor 488 marker (Figure 1, green). Thus, this assay is designed to show low colocalization if resolution is sufficient and clustering is low. For unexpanded small platelets, however, low image resolution leads to a linear relationship in the intensity scatter plot for the two color channels (Figure 1B). Such largely overlapping color signals only provide a meaningless high degree of – potentially false-positive – co-localization. In contrast, with 4x and 10x expanded platelets this linear relationship vanishes and colocalization analysis becomes discernible for different scenarios (Figure 1 C, D). Unfortunately, the resolution gain is accompanied by a loss in contrast due to reduced signal-to-noise ratio (Figure 1E). Moreover, the signal retention varies for different fluorescent dyes. While both antibodies were decorated with a similar number of fluorophores (see supplemental Table I), the signal retention is different for each color (Figure 1F, lower signal retention in the green channel than in the magenta channel). Low contrast and different degree of receptor visibility between channels require a careful preselection of the colocalization modality.

**Figure 1.**
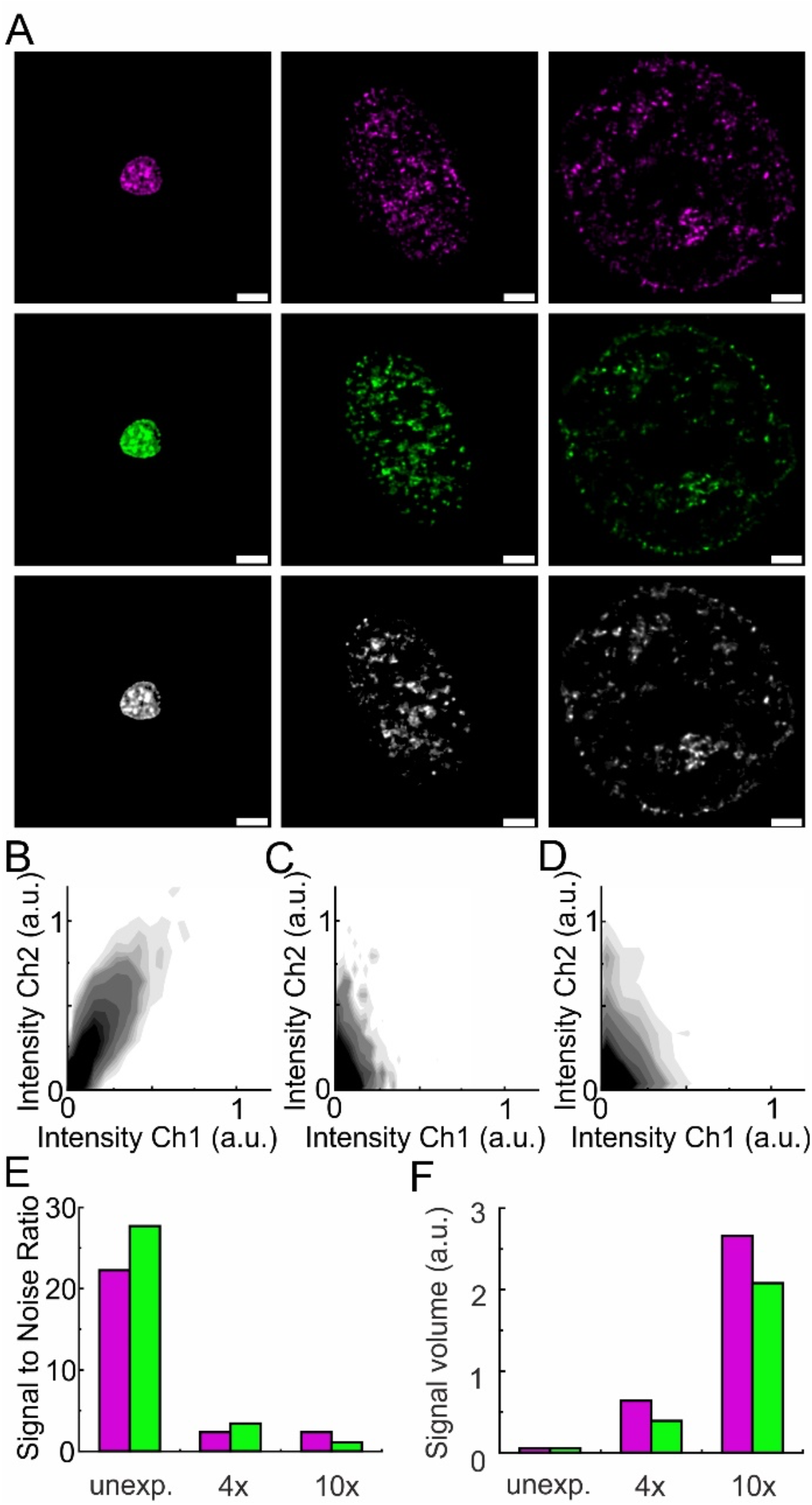
Colocalization analysis of receptor distributions on expanded platelets. **A**, Deconvolved confocal image planes of the GPIIb/IIIa receptor distribution in unexpanded (left panels), 4x expanded (middle panels) and 10x expanded (right panels) resting platelets immunolabeled with the monoclonal antibody MWReg30 (magenta: Alexa Fluor 594, green: Alexa Fluor 488, white: merged). **B-D**, Density plots of the signal intensities in channel 1 (Ch1, MWReg30-Alexa Fluor 594) and channel 2 (Ch2, MWReg30-Alexa Fluor 488) in logarithmic scale **(B)** unexpanded, **(C)** 4x expanded and **(D)** 10x expanded. **(E-F)** Signal to noise ratio **(E)** and signal volume **(F)** of the respective color channels derived from raw images in **(A)**. Scale bars 3 µm.

The Manders’ colocalization approach turned out to be the only applicable here as it is most crucial that the channel with fewer events still reports its overlapping fraction regardless of the degree of asymmetric signal loss in both channels. As the Manders’ colocalization coefficient, however, is less robust against noise than the commonly used Pearson’s colocalization coefficient, image preprocessing is required to exclude background signal efficiently.^13^ Along these requirements, we compiled an image processing pipeline providing a chromatic shift correction (Figure 2A) before a deconvolution step (Figure 2B) followed by background filtering (Figure 2 C,D). In the last step signal events are filtered by intensity threshold and size (detailed description in the supplementary methods).

**Figure 2.**
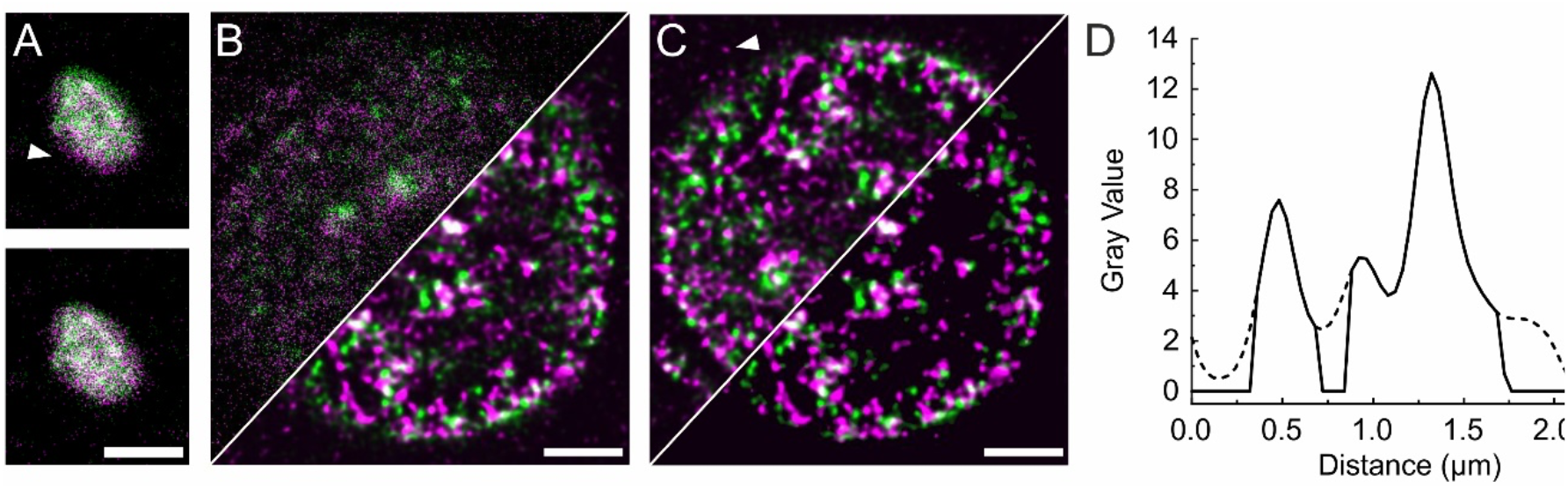
Image preprocessing. Prior to the colocalization analysis confocal z-stacks of GPIIb/IIIa receptor distributions (MWReg30; green: Alexa Fluor 488, magenta: Alexa Fluor 594) are processed by **(A)** a chromatic shift correction: raw unexpanded platelet, **(B)** image deconvolution: raw (left) and deconvolved (right) image of 4x expanded platelet, **(C)** masking the surface: deconvolved (left) and masked surface (right) of 4x expanded platelet. **(D)** Line plots of signal intensity before (dashed line) and after creating a masked surface (solid line) derived from images in **(C)**. Scale bars 3µm.

### Receptor distributions revealed by expansion of platelets

ExM increases the optical resolution level to the nanometer range; the exact resolution gain depends on the hydrogel composition and the respective expansion factor. This means that in principle a lateral resolution of down to 25 nm can be realized by 10x expansion and confocal microscopy; however, as this resolution gain comes at the expense of image brightness and contrast, we carefully dissected the resolution level that is required for colocalization analysis of the platelet receptors. First, we used realistic simulations (see Figure 3 and 4) to understand how the image quality and specificity would manifest itself under real experimental conditions. Therefore, randomly distributed single GPIIb/IIIa receptors or receptor clusters as fluorescent objects were computationally placed on a spheroid surface reflecting typical mouse platelet dimensions (Figure 3). From such platelet-like objects (Figure 3A) fluorescence images (Figure 3B-D representing maximal (left) and minimal (right) colocalization) were created including realistic background signal and noise (unexpanded: Figure 3B, 4x expanded: Figure 3C, 10x expanded: Figure 3D). The resulting signal-to-noise ratios of these simulated 3D images for all three expansion levels (none, 4x, 10x) and the two colocalization cases in comparison are plotted in dependence of increasing marker retention ratio. Even if the signal-to-noise level is higher for lower expansion factors, the colocalization coefficient becomes meaningless when looking at platelet receptor distributions with varying densities (Figure 4A) for a high and low colocalization test case (Figure 4B). For the high colocalization case, two markers can bind to one individual receptor at the same time giving rise to a high colocalization coefficient, while in the low colocalization case the two markers have less chance to colocalize as each receptor can only by occupied by either one marker (Figure 4B). In unexpanded platelets it is impossible to distinguish both cases reliably while this was possible for expanded platelets in the whole receptor density range. For a receptor density in the range ~500 /µm^2^ as we found for GPIIb/IIIa (see supplemental Figure III) 4x expansion is expected to only show a 10% drop in colocalization between the high and low case, while at 10x expansion a drop of 50% can be expected (Figure 4A). Thus, only 10x expansion provides sufficient dynamic range to pinpoint the differences of receptor colocalization in the two test cases. This advantage vanishes only if marker retention would drop below 50% (Figure 4C). Note that for both, 4x and 10x expansion, the achieved super-resolution is just not sufficient to resolve single molecules and thus may still leave a certain risk to detect “false-positive” colocalizations. Here in this case, even for very low receptor densities a Manders’ colocalization coefficient of 0.6 or 0.2, respectively, represents the low colocalization case. To understand how the loss of markers during homogenization and expansion influence these results, we additionally accounted for the clustered arrangement of GPIIb/IIIa on resting platelets as observed by single molecule localization microscopy (Figure 4D & supplemental Figure III). Decreasing marker retention in both channels means smaller colocalization values (Figure 4E), particularly in 4x expansion where the dynamic range between high and low colocalization is decreased for retention ratios below 50%; in contrast, for 10x expansion the performance is best maintained even beyond 50%. However, with further decreasing (asymmetric) retention ratio the analysis becomes prone to artifacts (Figure 3 & Figure 4F): When the retention ratio is fixed at 60% in channel A and the retention ratio of channel B ranges from 100% to 20% the results seem to be robust down to only 40% retention in channel B for both 4x and 10x expansion. In 4x expansion the dynamic range between the high and low colocalization are indistinguishable (high and low, Figure 4B) at 20% retention in channel B. The high colocalization is experimentally realized with two antibodies carrying either dye molecule or targeting different epitopes on GPIIb/IIIa. Thus, they can bind to a single receptor at the same time (supplemental Figure IV, case I).

**Figure 3.**
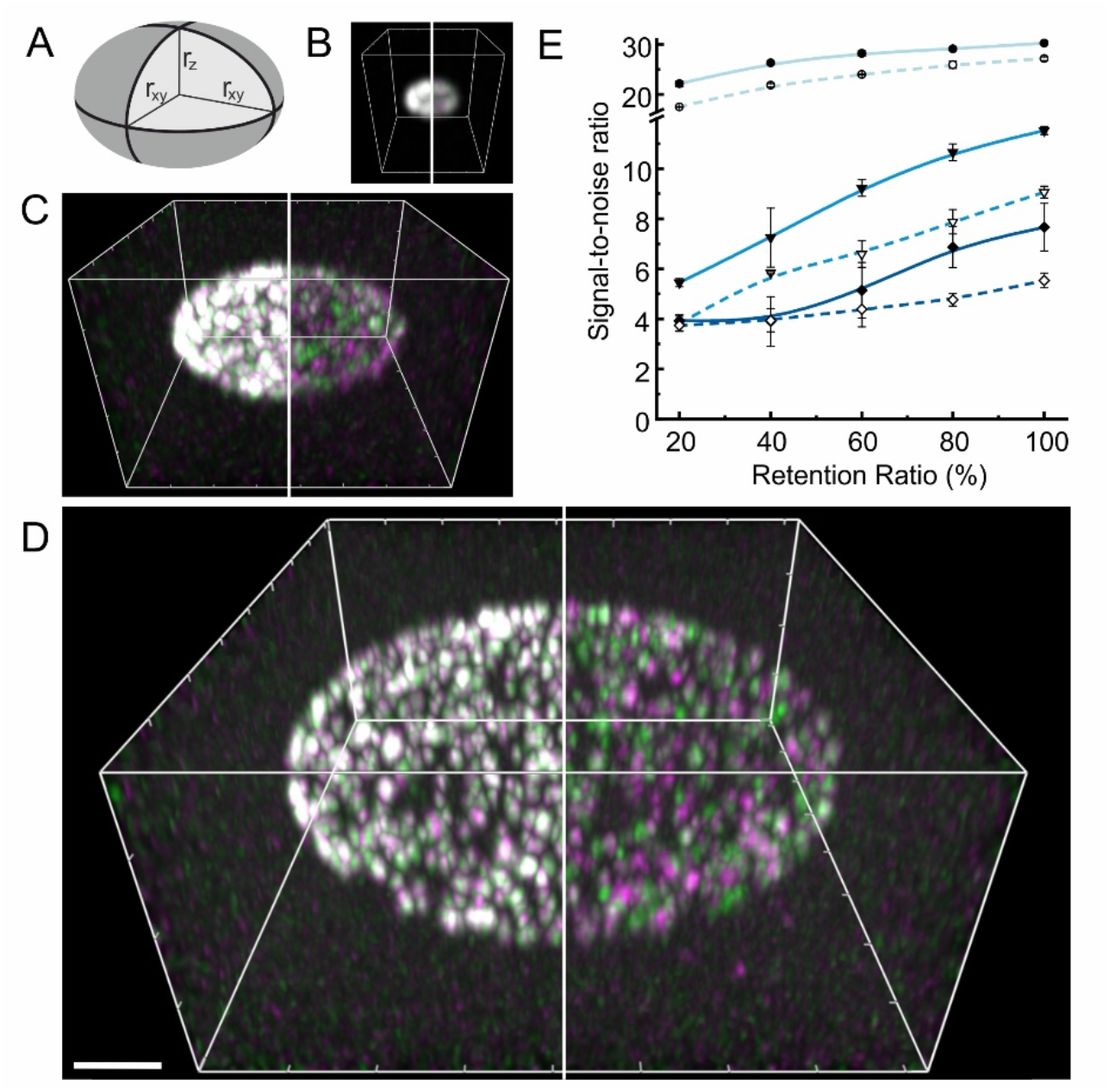
Simulated platelet receptor distributions. **A**, Randomly distributed single GPIIb/IIIa receptors or receptor clusters positions were placed on a spheroid surface reflecting typical mouse platelet dimensions (r_xy_ = 1.5 µm and r_z_ = 0.5 µm). **B-D**, Based on the simulated receptor distributions and the antibody and marker combination (Case I: left, Case II: right) realistic 3D fluorescence images were created including the generation of background signal and noise for **B** unexpanded, **C** 4x expanded (brightness increased 2x) and **D** 10x expanded platelets (brightness increased 10x). In the images presented here the marker retention ratio is 100%. **E**, Signal-to-noise ratio of simulated 3D images in Case I (solid lines, filled symbols) and Case II (dashed lines, open symbols) of unexpanded (light blue), 4x expanded (medium blue) and 10x expanded platelets for a marker retention ratio ranging from 20% to 100%. Scale bar B-D 4 µm

**Figure 4.**
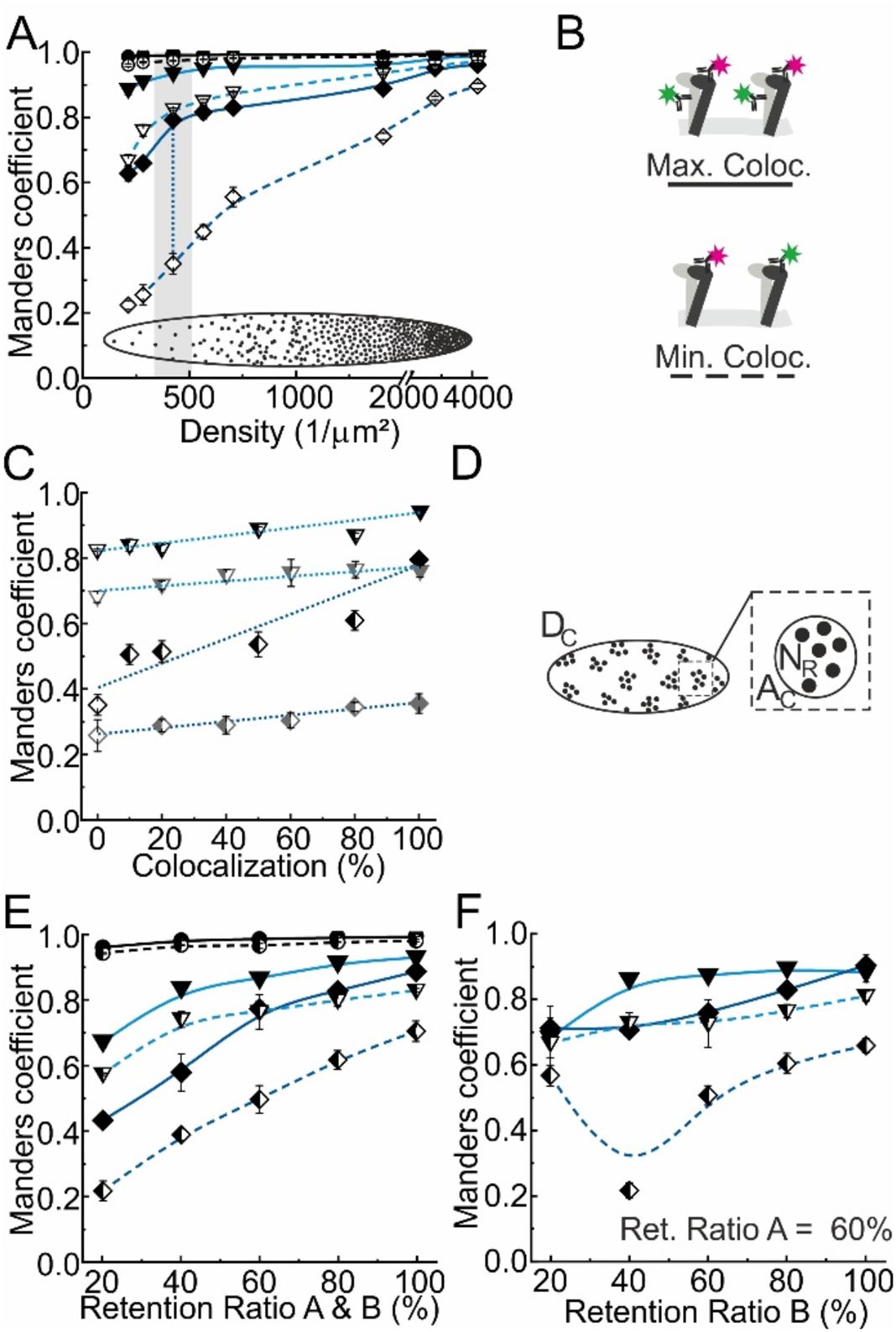
Investigating the influence of receptor densities and label retention ratios of the accuracy and resolution of colocalization analysis with ExM based on simulated receptor distributions. **A**, Density dependence of the Manders’ coefficients for the maximum (solid lines & filled symbols) and minimum (dashed lines & open symbols) colocalization cases illustrated in (B); non-interacting receptor distributions on unexpanded (black), 4x expanded (light blue) and 10x expanded platelets (dark blue) with corresponding SD (error bars). The lines show the trend with a B-spline, the gray shaded area highlights the expected range of the average density of GPIIb/IIIa receptors and the dotted line indicates the dynamic range of the Manders’ coefficient. **B**, Cartoon of the maximum and minimum colocalization test case. **C**, Manders’ coefficients with corresponding SD (error bars) of simulated receptor distributions with different degrees of colocalization on 4x (triangles) and 10x (diamonds) expanded platelets for a label retention ratio of 100% (black symbols) and 50 % (gray symbols) with a linear fit (dotted lines) at the receptor density expected for GPIIb/IIIa (dotted line in in A). **D**, Cartoon of a clustered receptor distribution indicating the cluster density D_C_, cluster area A_C_ and the number of receptors per cluster N_R_ (for GPIIb/IIIa: D_C_=70 µm^-1^, A_C_=300 nm and N_R_=7, see supplemental Figure III). **E**,**F** Manders’ coefficients with corresponding SD (error bars) in simulated receptor cluster distributions for the maximum (solid lines, filled symbols) and minimum (dashed lines, half-filled symbols) colocalization test cases in the case of **(E)** equal and **(F)** asymmetric scenarios of label retention, unexpanded (black), 4x expanded (light blue) and 10x expanded platelets (dark blue).

Next, we would like to look at true experimental data. For image preprocessing and analysis, it is important to include only platelets above the largest cutoff and with suitable orientations to avoid bias in the subsequent colocalization analysis (Figure 5). Thus, we excluded not fully expanded platelets from further analysis (Figure 5A), and made sure that subpopulations of platelets with different orientations are correctly reflected in the distribution of Manders’ colocalization coefficients (Figure 5 B,C). Now, we are fully prepared to interrogate three different colocalization scenarios: For the high colocalization case two antibodies targeting two different epitopes of GPIIb/IIIa are used carrying Alexa Fluor 488 and Alexa Fluor 594 respectively (Figure 6, case I). For the low colocalization, only one antibody targeting a single epitope of GPIIb/IIIa is used, carrying either Alexa Fluor 488 or Alexa Fluor 594 (Figure 6A, case II). As an additional case (case III, Figure 6) we introduce two independent receptors, GPIIb/IIIa and GPIX, which are targeted with distinct fluorophores and no colocalization is expected.

**Figure 5.**
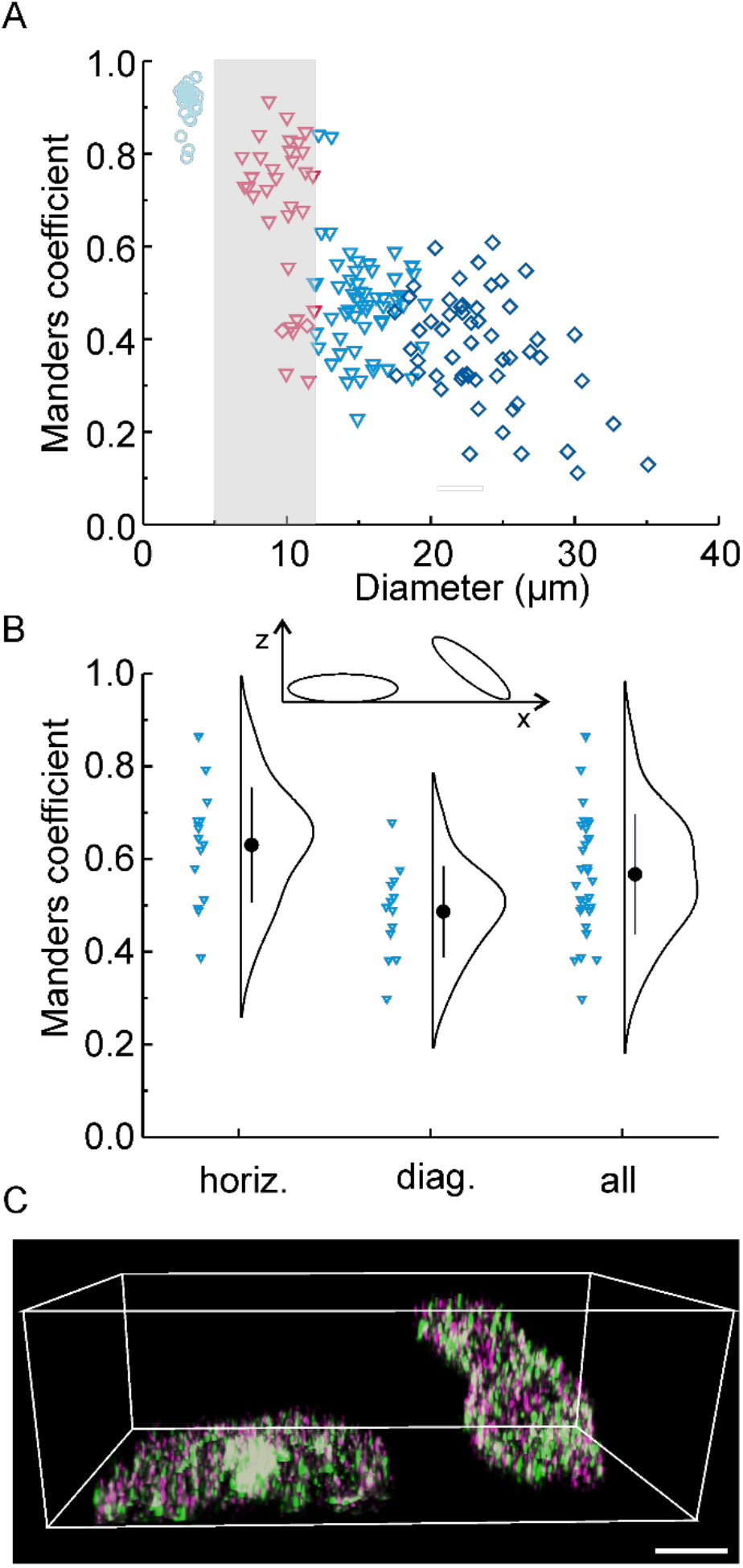
Manders’ colocalization coefficient dependence on expansion factor and orientation. **A**, Manders’ colocalization coefficient vs. diameter of unexpanded (circles), 4x expanded (triangles) and 10x expanded (diamonds) resting platelets labeled with anti-GPIIb/IIIa antibodies (MWReg30) carrying Alexa Fluor 488 and anti-GPIX antibodies (p0p6) carrying Alexa Fluor 594 (Case III). Expanded platelets with a diameter below 12 µm were excluded from further analysis (marked in red) (**B-C**) **B**, Distributions of the Manders’ colocalization coefficients of 4x expanded resting platelets labeled with an anti-GPIIb/IIIa antibody (MWReg30) carrying either Alexa Fluor 488 (green in C) or Alexa Fluor 594 (magenta in C, Case II) selected for horizontal (horiz.) or diagonal (diag.) orientation and without any selection (all).**C**, Two example platelets from the data in B exhibiting horizontal (left) and diagonal (right) orientation. Scale bar 4 µm

**Figure 6.**
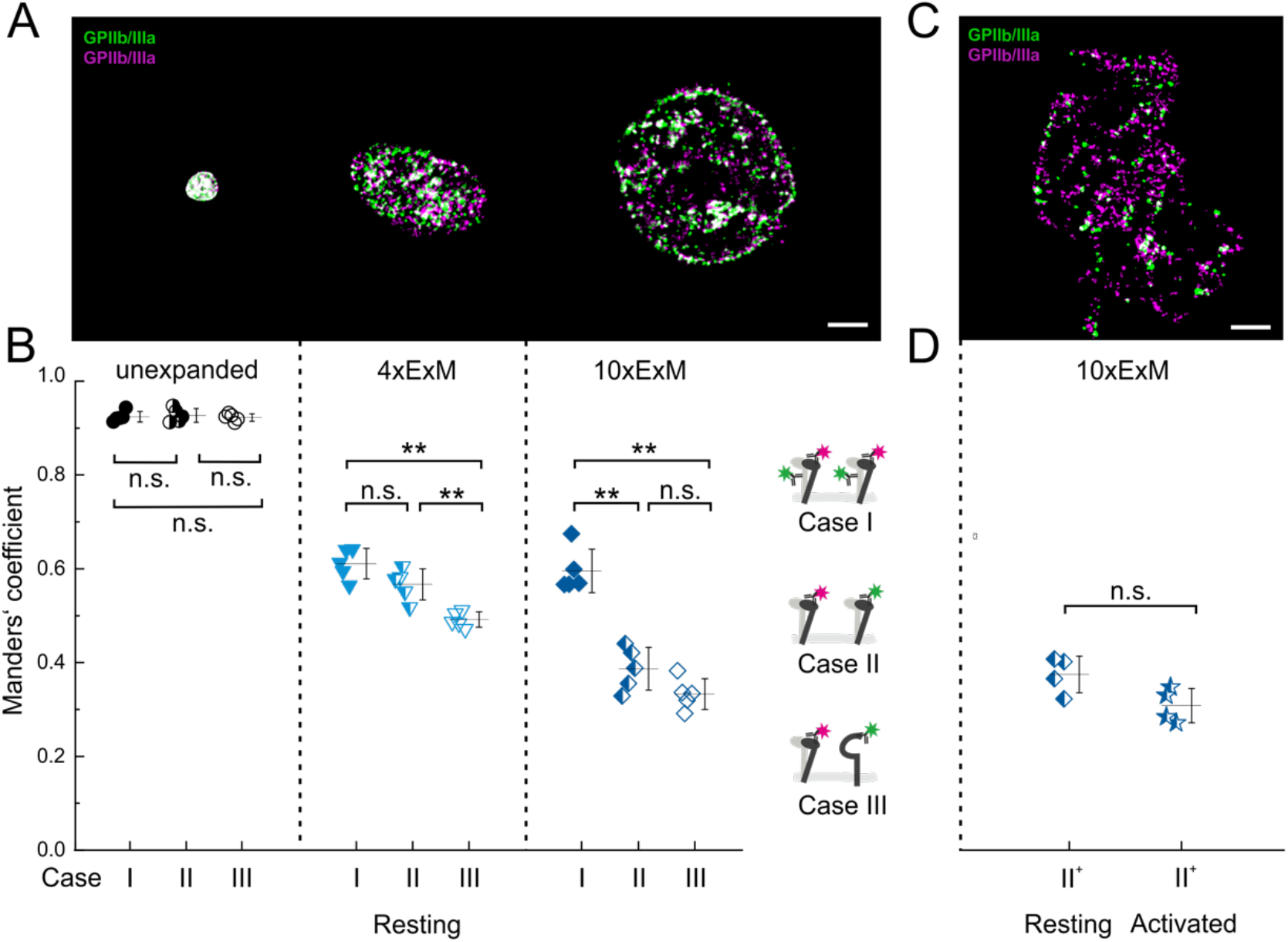
Performance of ExM based colocalization analysis. **A**, Deconvolved confocal image slices of GPIIb/IIIa receptors on unexpanded (left), 4x expanded (middle) and 10x expanded (right) resting platelets,all directly labeled with fluorescent antibodies (Alexa Fluor 488 (green) or Alexa Fluor 594 (magenta). **B**, Comparing the distributions of the average Manders’ coefficients of groups of six with corresponding SD (error bars) of unexpanded (left, black circles), 4x expanded (middle, light blue triangles) and 10x expanded (right, dark blue diamonds) resting platelets labeled with three different combinations of antibodies (Case I: filled symbols, Case II: half-filled symbols & Case III: open symbols). **C**, Deconvolved confocal image slice of GPIIb/IIIa receptors on a 10x expanded activated platelet, directly labeled with fluorescent antibodies (Alexa Fluor 488 (green) or Alexa Fluor 594 (magenta).**D**, Distributions of the average Manders’ coefficients of groups of five with corresponding SD (error bars) of 10x expanded resting (dark blue diamonds) and activated platelets (dark blue stars) with labeling scheme Case II^+^ (Antibody batch 2 with lower density of labeling, see supplemental Table I). n.s.: p > 0.05, *: p ≤ 0.05, **: p ≤ 0.01, (Kruskal-Wallis ANOVA test), scale bars 5 µm.

For unexpanded platelets (Figure 6A left) high Manders’ colocalization coefficients are observed for all three test cases and differences in the marker arrangement are not resolved (Figure 6B). In line with previous reports by others^30^ our results here underscore that conventional confocal microscopy completely fails when it comes colocalization studies of densely packed targets. By expanding the platelets 4x the coefficient for case I drops down to ~0.6 due to the limited retention ratio. It can already be distinguished from case III with a dynamic range of 0.15 between maximum and minimum colocalization while the resolution is insufficient to distinguish it from case II. The increase in resolution achieved with 10x expansion allows to extend the dynamic range of the Manders’ colocalization coefficient to 0.25. While in the 4x expansion experiments the limited resolution still leads to an apparent Manders’ colocalization coefficient of ~0.5 for case II and III, it drops down to ~30% for 10x expansion with a clear separation between the high and low colocalization. Simulated data shows 4x expansion does not provide the resolution required at the given receptor densities. Interestingly, case II yields a higher, but not significantly different colocalization coefficient than case III, maybe due to the higher abundance of the GPIIb/IIIa receptors. Fortunately, a neighbor-distance analysis of 10x expanded platelets allows to distinguish colocalized, clustered and non-interacting receptor distributions (supplemental Figure V) overcoming its lack of molecular resolution.

In an independent experiment we compared the GPIIb/IIIa distribution on resting and activated platelets to assess whether platelet activation affects the formation of integrin clusters. Surprisingly, GPIIb/IIIa colocalization coefficients of activated platelets were comparable to those in resting platelets (Figure 6C,D), suggesting that the organization of GPIIb/IIIa receptors is conserved during platelet activation.

## Discussion

This study shows that the combination of ExM and confocal imaging is a powerful tool to study platelet receptor organization and interaction in small blood platelets and dense and highly abundant receptor scenarios where other high- and super-resolution methods fail. Conventional confocal microscopy alone is not suited as demonstrated here. Based on an optimized sample preparation and tailored image preprocessing and analysis procedure we developed a colocalization analysis approach which allows to classify the distribution and interaction of high-density platelet receptor populations. Our findings suggest, that GPIIb/IIIa receptors are present in pre-formed clusters already on resting blood platelets. These pre-formed clusters might play a role in enhancing platelet adhesion and aggregation following inside-out activation and thereby contribute to the rapid platelet responses upon vascular injury. The unexpected observation that these clusters are not affected by platelet activation (Figure 6D) supports this hypothesis. These data indicate that – at least in the absence of an immobilized ligand – no significant rearrangement of GPIIb/IIIa clusters relative to each other is observed. One might speculate that these uniformly distributed clusters represent preassembled outside-in signalosomes that are ‘ ready to go’ upon integrin-ligand binding or inside-out signaling. Thereby, they enable ligand binding on the entire platelet surface, allowing a rapid response to external stimuli. Clearly, further studies are needed to elucidate the receptor interplay on the platelet surface. In particular, the effects of immobilized ligands on the distribution of GPIIb/IIIa clusters are worth studying, to investigate whether such ligands would result in bigger GPIIb/IIIa clusters along the ligands similar to GPVI.^31,32^On the other hand it is also possible that these uniformly GPIIb/IIIa clusters are critical for the mechanosensing capacity of platelets and potentially platelet migration.^33,34^

In any case our study successfully addresses the special requirements posed by the small platelet size and high receptor density. While alternative super-resolution microscopy techniques can provide single molecule resolution, which is not achieved here, the presented approach stands out by its accessibility as it does not require an advanced super-resolution setup. Furthermore, the combination with confocal microscopy allows for the implementation of both, 3D and multicolor imaging, still bottlenecks for many super-resolution techniques.

However, regarding colocalization analysis the approach itself holds intrinsic challenges as it not only lacks the mentioned single molecule sensitivity, but also suffers from a significant loss of fluorescent markers. We evaluated the performance of ExM based colocalization analysis regarding receptor density and retention ratios based on simulated GPIIb/IIIa receptor distributions. While an expansion of 4x still provides robust results at retention ratios below 50%, the dynamic range between high and low colocalization is significantly lower than for 10x expansion and especially for receptor densities above 1000/µm^2^ only 10x expansion provides sufficient resolution to distinguish both cases. As mentioned before, the marker retention ratio greatly depends on the sample processing as well as the molecular properties of the fluorescent dye. For the organic dyes Alexa Fluor 488 and Alexa Fluor 594 used in this study, retention ratios of ~50% have been reported.^19^

The higher resolution achieved by 10x expansion comes at the cost of signal dilution, a longer and harsher sample preparation and a decreased mechanical stability of the hydrogel. To address the issue of signal loss and dilution, several modifications of the anchoring chemistry and homogenization procedure have been reported to allow post expansion labeling.^35–37^

Despite these limitations ExM will help to improve our understanding of platelet receptor distribution. This will have implications on our understanding of platelet integrin receptor stoichiometry and spatial organization like in the present study. We have established ExM for murine platelets as this allows to combine this attractive technology with transgenic models to investigate further research questions in the platelet field, like the existence/relevance of clusters of (hem)ITAM receptors,^38,39^ lipid rafts^40^ or the question whether different pools of α-granules exist in platelets.^41,42^ By combining ExM with advanced imaging techniques it is even possible to achieve molecular resolution^37,43–45^ if the marker and labeling concept allows. Thus, we believe that ExM for platelet imaging has a bright future and may soon help to decipher complex receptor interactions within the nanometer range.

## Acknowledgments

K.G.H., D.S., B.N. and H.S.H conceived and designed the experiments. H.S.H. performed the simulations for the resting platelets and supervised the ExM experiments. M.A. developed the platelet expansion protocol and the image acquisition procedure and developed the image processing and colocalization analysis procedures. M.A. and S.M. performed the ExM experiments and analysis on resting platelets. P.G. performed the ExM experiments on activated platelets while L.M.C.E. performed the data analysis. C.K. and M.M. performed the dSTORM imaging and cluster analysis. P.G. and S.M. performed the confocal imaging on activated and resting platelets respectively. V.G. screened the antibody incubation times for live and post-fix labelling and assisted with the animal handling. D.S., B.N. and V.G. advised with the platelet preparation and immunolabeling.

B.N. provided the platelet antibodies. K.G.H., H.S.H and D.S. wrote the paper. All authors read and approved the final manuscript. We thank Shazeb Ahmad and Christoph Kempe for assistance in protocol development of ExM, Dr. Katharina Remer for support with documentation and management of animal experimentation and welfare, as well as Harald Schulze and Markus Sauer for fruitful discussion around platelet biology, expansion microscopy and colocalization analysis. Thanks to A. Balakrishnan for proof-reading.

