## Supplementary Material for "Mapping densely packed GPIIb/IIIa receptors in murine blood platelets with expansion microscopy"

* corresponding authors:

**Other supplementary materials for this manuscript include the following:**

Zenodo Repository: https://doi.org/10.5281/zenodo.4117555

#### Online Supplementary Methods

**Colocalization analysis.** For colocalization, single platelets were cropped in Imaris 9.4.0 (BITPLANE, Oxford Instruments), background was excluded using intensity and volume thresholded surfaces and analyzed with Imaris Coloc module. The Manders coefficient of the channel with the lower receptor density (lower total surface volume) was taken into account.

**Statistical Analysis.** Data were tested for normality (Shapiro-Wilk) and statistical analysis between the groups was performed using a nonparametric test (Kruskal–Wallis ANOVA). Data is expressed as mean ± SD, and $P< 0.05$ was considered statistically significant.

**Receptor density estimation.** To estimate the number of receptors per platelet, a workflow was developed to count the number of events per surface area in one platelet. All following steps were performed using Imaris 9.2.1 (BITPLANE, Oxford Instruments). The number of receptors on the platelet surface was counted by creating surface objects for each event. The receptor events were identified in the surface module with the settings “smooth”, “background subtraction: local contrast, 0.4 µm”, “threshold: 10.6”, “split touching objects: 0.2 µm”, “seed point filters: mean intensity > 20” and “classify surfaces filters: number of voxels > 40”. The platelets surface area and volume was estimated, using the Imaris Cell Module with the settings “detect cell only”, “detect cell boundary form cytoplasm”, a smoothing of 0.5-5 µm. The intensity threshold was adjusted to cover the whole platelet with a closed surface. Since some platelets were cut off at the bottom during imaging, the “missing” area was calculated and subtracted from the surface area as follows: All voxels in the previously modulated cell were masked to intensity 100 in a new channel. A surface with a thickness of 1 voxel thick was created with the surface module at the cut off and half of the created surface area was subtracted from the total platelets surface area.

**Simulation:** The image stacks of simulated platelet receptor distributions were generated with MatLab (Mathworks) based on evenly distributed positions on a spheroid surface with the original extension of $r_z$=0.5 µm and $r_{xy}$=1.5 µm. Based on the coordinate positions of randomly picked receptor positions and taking into account the retention ratio R a three dimensional image stack was reconstructed with a PSF size of $PSF_{x,y}$=200 nm and $PSF_z$=500 nm, a photon count of 1000 per event, a background level of 5 counts, a signal-to-noise ratio of 5, Poisson distributed noise and a pixel size of x,y=40 nm and z=180 nm. For an expansion factor of 4 and 10 the coordinate positions were extended accordingly. The simulated image stacks were deconvolved and analyzed as described in the previous sections.

**dSTORM sample preparation:** 8-well chamber slides (Cellvis) were coated with 200 µL 2M glycine for 10 min. at RT for resting platelets. Platelets were washed as described elsewhere.[1] Briefly, mice were anesthetized using isoflurane and bled into 300 µl heparin (20 U/ml in TBS, Ratiopharm). The blood was centrifuged twice at 300 *g* for 6 min to obtain platelet-rich plasma (PRP). PRP was supplemented with 0.02 U/ml apyrase (A610, Sigma-Aldrich) and 0.1 µg/ml PGI2 (P6188, Sigma-Aldrich) and platelets were pelleted by centrifugation at 800 g for 5 min, washed twice with Tyrodes-HEPES buffer (134 mM NaCl, 0.34 mM Na2HPO4, 2.9 mM KCl, 12 mM $NaHCO_3$, 5 mM HEPES, 5 mM glucose, 0.35% BSA, pH 7.4) containing 0.02 U/ml apyrase and 0.1 µg/ml $PGI_2$. The platelets were allowed to rest for 30 minutes prior to experiments and platelet counts were adjusted to 300,000 per µL in Tyrode's buffer without $Ca^{2+}$. 60 µL of the platelet suspension was added to 200 µL Tyrode's buffer without $Ca^{2+}$ in each glycine-coated well and kept at 37°C for 30 min. as well. After 30 min. The supernatant was discarded, and the platelets were fixed with 200 µL prewarmed glyoxal-solution and centrifuged at 300 *g* for 5 min at RT. Subsequently the samples were quenched with 0.1% $NaBH_4$ for 15 min. and blocked with 5% BSA for 1h. Staining was done for 30 min. at 37°C with 1 µg Alexa Fluor 647 coupled anti-αIIbβ3 (GPIIb/IIIa) antibodies MWReg30 and JON6 (generated and modified in our laboratory as described previously[2,3]) in 300 µL PBS in each well.

**dSTORM microscopy:** Two-color dSTORM was performed on a widefield setup based on an inverted microscope (Olympus IX-71) equipped with an oil immersion objective (60x, NA 1.45; Olympus). Alexa Fluor 647 and Alexa Fluor 532 were excited with the appropriate laser systems (Genesis MX639 and MX514, Coherent) at irradiation intensities between 2-10 kW/cm$^2$. Emission light was separated from the excitation light using a dichroic mirror (ZT 405/514/635rpc, Chroma Technology Corp.), spectrally filtered by a bandpass filter (Em01-R442/514/647-25; Semrock), transmitted by a dichroic beam splitter (ZT 405/514/635rpc, Chroma Technology Corp.) and respective emission filters (Brightline HC 679/41, Brightline HC 582/75, Semrock), before being projected on two EMCCD camera chips (iXon Ultra DU-897, Andor). Imaging was done with epifluorescent (EPI) illumination and an exposure time of 20 ms for 15,000 frames (red dye) and 30,000 frames (green dye) in photoswitching buffer containing 100 mM β-mercaptoethylamin, pH 6.9, without oxygen scavenger as previously described[4]. Fluorescent beads (100 nm TetraSpeck beads, T7279, Thermofisher) were imaged in both channels over the whole ROI to later correct for chromatic aberrations. dSTORM images were reconstructed using the open source software rapidSTORM 3.3,[5] matrices for chromatic aberration corrections were done with the bUnwarpJ[6] plugin in Fiji.[7] The cluster analysis was performed with a custom written Python-based script.

105 **Supplementary Figures**

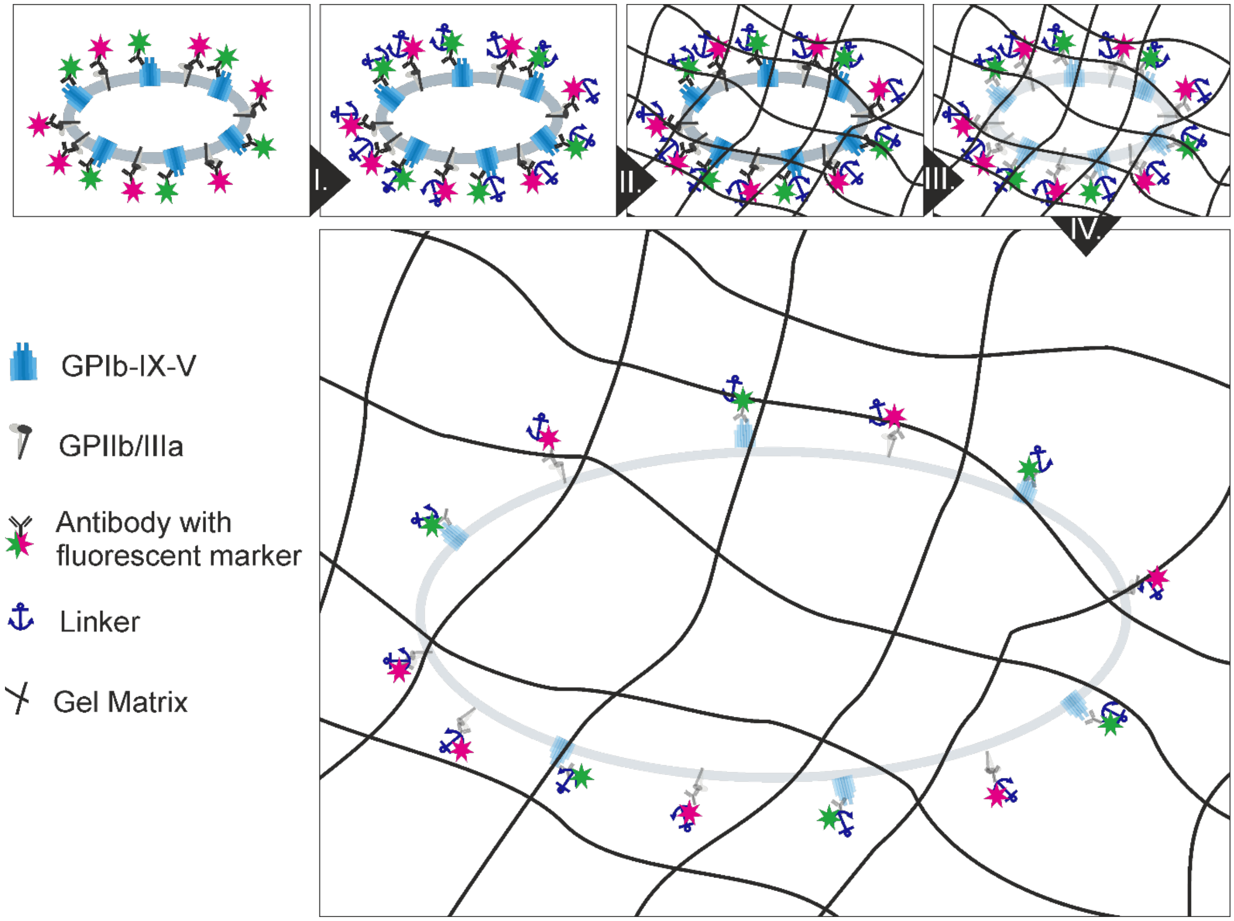


**Supplementary Figure I. The principle of expansion microscopy**.

After labeling the proteins of interest with fluorescent markers via antibodies (I. Anchoring) a linker reagent is added that binds to proteins and is able to covalently anchor them into the gel matrix. (II. Polymerization) The sample is immersed in a monomer solution and the polymerization process is started by adding an initiator reagent. (III. Digestion) To ensure isotropic expansion the sample is homogenized by protease digestion. (IV. Expansion) Finally the sample is expanded to 4x or 10x its original size in $ddH_2O$.

106

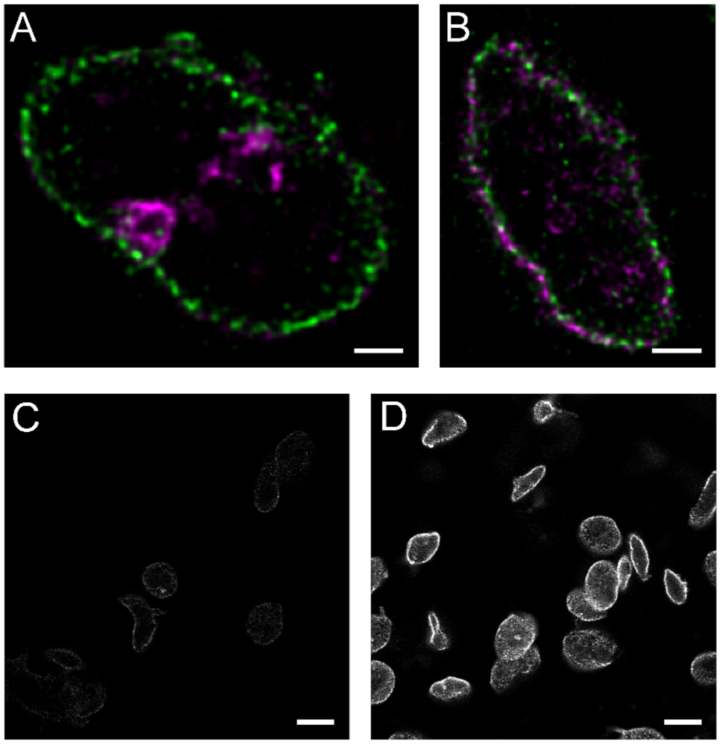


**Supplementary Figure II: Optimized sample labeling and fixation procedure.**

**A,B** Live and post-fix staining: Deconvolved confocal images of 4x expanded resting platelets labeled with anti-GPIIb/IIIa antibody carrying Alexa Fluor 488 (MWReg30, magenta) and anti-GPIX antibody (p0p6, green) carrying Alexa Fluor 594 before (**A**, live) and after fixation with glyoxal (**B**, post fix). **C,D** Sample fixation with PFA and glyoxal: Deconvolved confocal images of resting platelets fixed with PFA **(C)** and glyoxal **(D)**, stained with an anti-GPIX antibody, p0p6, carrying Alexa Fluor 594 and expanded 4x. Scale bars A,B 2 μm and C,D 10 μm.

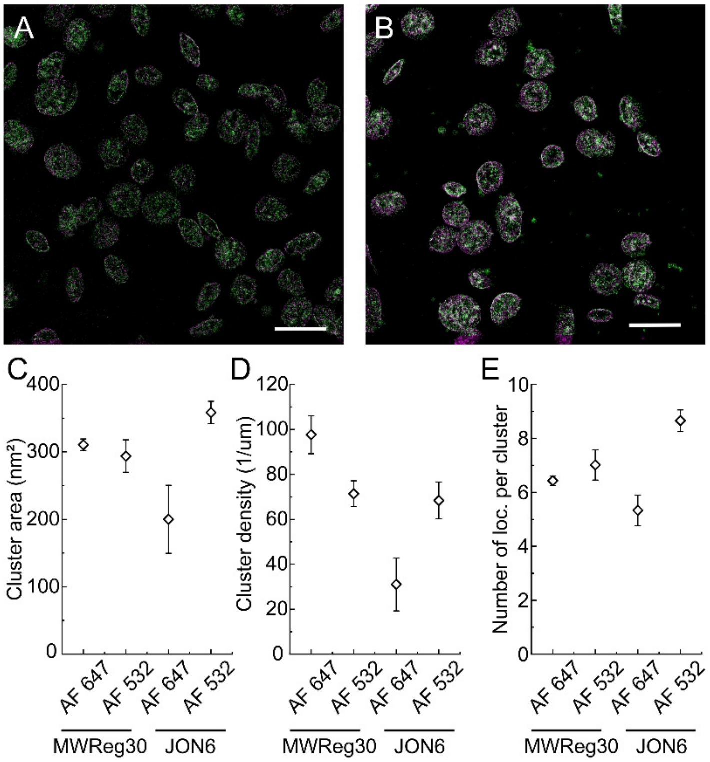


**Supplementary Figure III. GPIIb/IIIa cluster analysis based on single molecule localization microscopy.**

Super-resolved reconstruction of GPIIb/IIIa localizations acquired with dual-color *d*STORM of resting platelets labeled with two anti-GPIIb/IIIa antibodies which can bind to a single receptor at the same time. **A,** IgG JON6-Alexa Fluor 647 and IgG MWReg30-Alexa Fluor 532 and **B,** IgG JON6-Alexa Fluor 532 and IgG MWReg30-Alexa Fluor 647. **C-E,** Distribution of the cluster area (**C**), the number of localizations per cluster (**D**) and the cluster density (**E**) for the different antibody-dye combinations. Scale bars A, B 5 µm

108

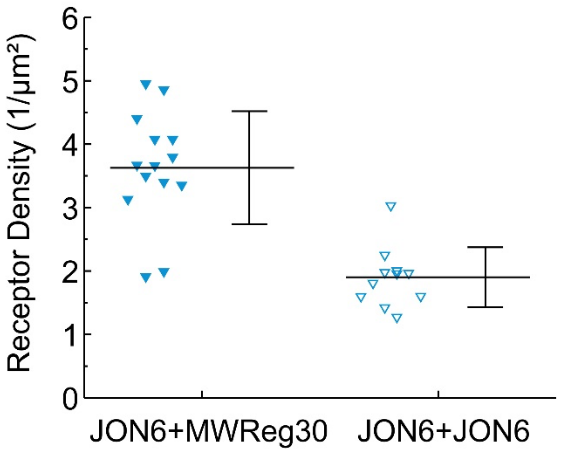


**Supplementary Figure IV. Simultaneous binding both anti-GPIIb/IIIa antibodies.**

Receptor density calculated based confocal images of 4x expanded resting platelets labeled with IgG MWReg30-Alexa Fluor 488 and IgG JON6-Alexa Fluor 546 (Case I, mean ± standard deviation: 3.6 ± 0.9) or with IgG JON6 carrying either Alexa Fluor 488 or Alexa Fluor 546 (Case II, mean ± standard deviation: 1.9 ± 0.5). In case I the receptor density is increased by a factor of ~1.9

109

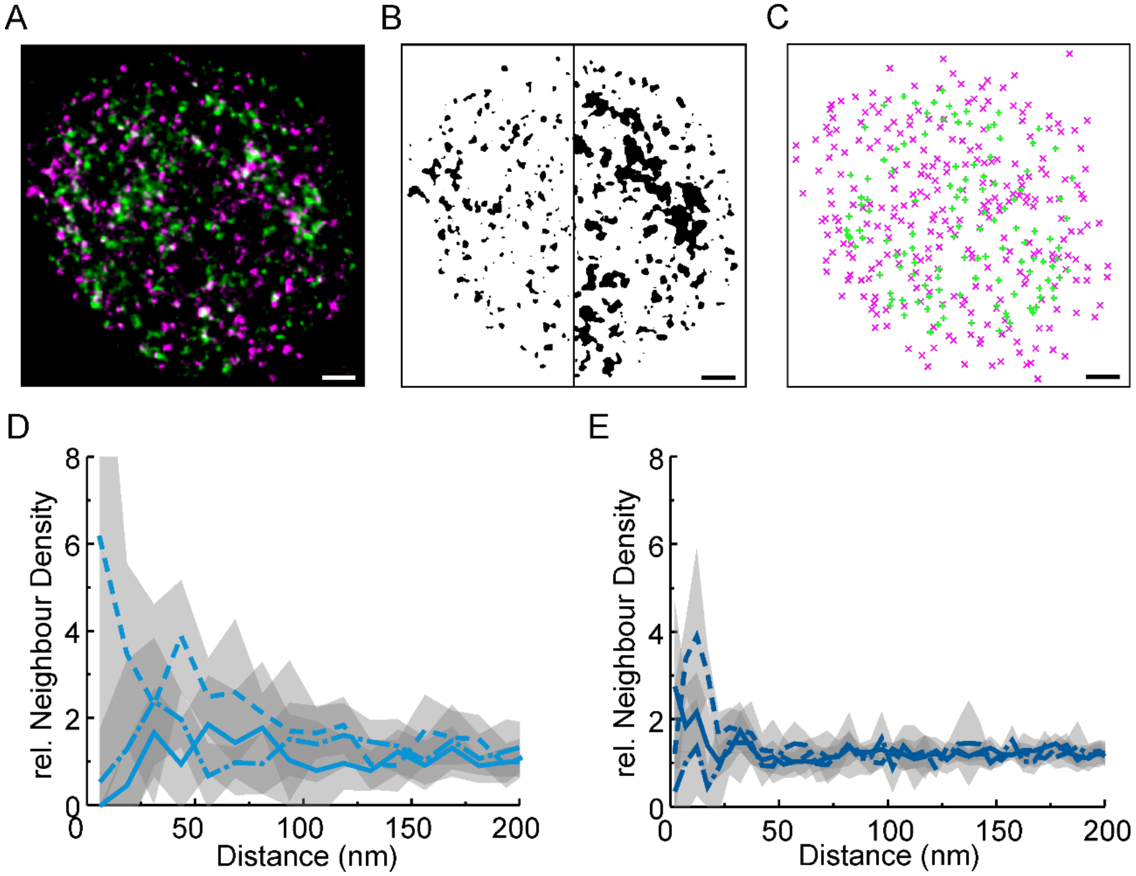


**Supplementary Figure V. ExM-based neighbor distance analysis.**

The receptor distribution in (**A**) a region of flat and plane membrane (here Case I: two antibodies carrying Alexa Fluor 488 (MWReg30, green channel) or Alexa Fluor 594 (JON6, magenta channel) can bind simultaneously on the same GPIIb/IIIa receptor) is analyzed by localizing receptors based (**B**) on "Otsu" thresholding followed by (**C**) mapping the center of mass of the detected objects. (**D-E**) The average object density detected in five images of (**D**) 4x and (**E**) 10x expanded platelets of Case I (solid line), Case II (dashed line, directly immunolabeled with a monoclonal antibody targeting GPIIb/IIIa (MWReg30) carrying either Alexa Fluor 594 or Alexa Fluor 488) and Case III (dash-dotted line, directly immunolabeld with monoclonal antibodies carrying Alexa Fluor 488 targeting GPIIb/IIIa (JON6) or carrying Alexa Fluor 594 targeting GPIX, p0p6). The corresponding SD is illustrated by the gray shaded area. Scale bars 2 µm.

Density mean ± SD 1/µm² :

Case I: 4xCh1 1.05 ± 0.40, 4xCh2 0.67 ± 0.17, 10xCh1 1.11 ± 0.18, 10xCh2 0.84 ± 0.15,

Case II: 4xCh1 0.90 ± 0.10, 4xCh2 0.65 ± 0.18, 10xCh1 0.76 ± 0.22, 10xCh2 0.67 ± 0.15,

Case III: 4xCh1 1.02 ± 0.25, 4xCh2 0.80 ± 0.26, 10xCh1 0.64 ± 0.32, 10xCh2 0.88 ± 0.15,

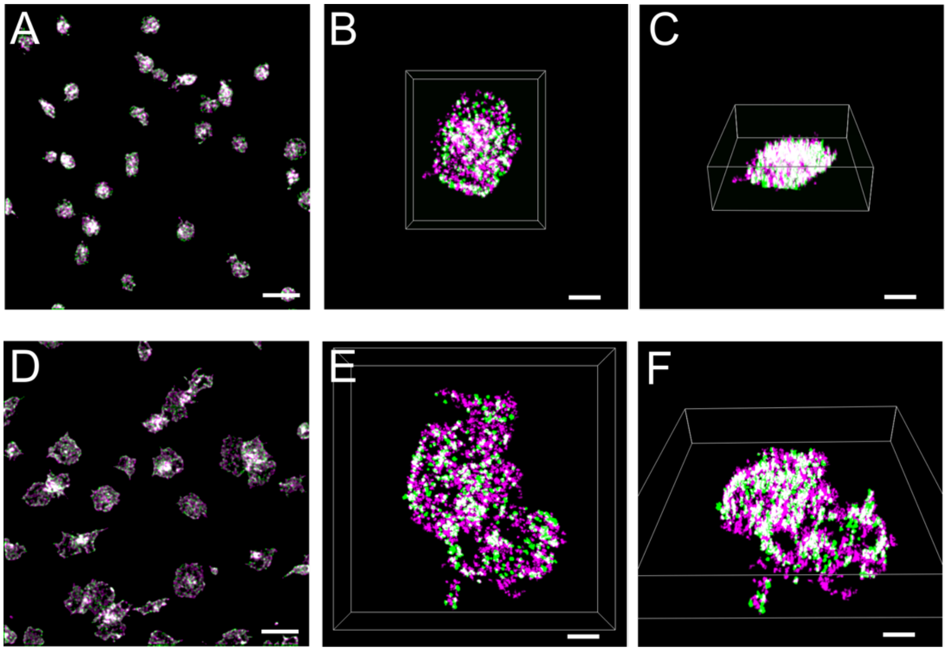


**Supplementary Figure VI. Expansion of resting and activated platelets.**
Deconvolved confocal images of resting (**A-C**) and activated platelets (**D-F**) labeled with anti-GPIIb/IIIa antibody (MWReg30, batch 2) carrying either Alexa Fluor 488 (green) or Alexa Fluor 594 (magenta). **A,** Single z-plane of unexpanded resting platelets settled on a glycine coated coverslip. Top view (**B**) and tilted view (**C**) of a 3D image rendering of a 10x expanded resting platelet. **D,** Single z-plane of unexpanded activated platelets settled on a glycine coated coverslip. Top view (**B**) and tilted view (**C**) of a 3D image rendering of a 10x expanded activated platelet. Scale bars 5 µm.

**Supplementary Table I: Degree of labeling of the antibodies**

| Receptor - Clone | Marker | Degree of labeling |
|---|---|---|
| GPIIb/IIIa – (Batch1) MWReg30 | Alexa Fluor 488 | 4.8 |
| | Alexa Fluor 594 | 3.4 |
| GPIIb/IIIa – (Batch 2) MWReg30 | Alexa Fluor 488 | 2.7 |
| | Alexa Fluor 594 | 3.3 |
| GPIIb/IIIa – JON6 | Alexa Fluor 488 | 3.3 |
| | Alexa Fluor 594 | 2.8 |
| GPIX – p0p6 | Alexa Fluor 594 | 6.4 |

#### Supplementary Protocol A: Stock solution recipes

##### I. Monomer Solutions

**Warning:** Sodium Acrylate is irritating to the eyes, to the respiratory system and the skin. Wear appropriate protective equipment and work under a fume hood.

**Purity note:** Sodium Acrylate sometimes comes with a variable purity level, which can affect performance. Best way to store it: solve whole bottle (38 g/100 ml (38wt%)). Sterilize through filter and aliquot. Store at -20°C in a desiccated environment. To ensure sufficient purity of the reagent discard batches that exhibit a **strong yellow tint** in solution.

**Warning:** DMAA has acute toxic effects on skin, on eyes and after oral contact. Wear appropriate protective equipment and work under a fume hood.

**Warning:** Acrylamide is **carcinogenic,** wear appropriate protective equipment and work under a fume hood. Any residues should be only discarded after complete polymerization.

**Keep everything cool during the whole process to prevent gelation, so work in a box with ice.**

###### Ia. 4x Monomer Solution

1. 50 g/100ml acrylamide solution: prepare stock solution with 5 g in 10 ml ddH2O and freeze it.
2. 2 g/100 ml N,N'-methylenebisacrylamide: prepare stock with 0.2 g in 10 ml ddH2O and freeze it.
3. 292 g/l sodium chloride: 1.46 g and fill up with ddH2O to 5ml.

| V [ml] | Reagents | Stock [g/100 ml] | Final [g/100 ml]=(w/w) |
|---|---|---|---|
| 2.27 | Sodium acrylate | 38 | 8.625 |
| 0.5 | Acrylamide | 50 | 2.5 |
| 0.75 | N,N'-Methylenebisacrylamide | 2 | 0.15 |
| 4 | Sodium chloride | 29.2 (0.5 M) | 11.69 (2 M) |
| 2.48 | PBS | 10x | 1x |
| 10 | **Final Volume** | | |

4. Fill reaction tubes with 480 µl of 4x monomer solution & store at -20°C.

**<u>Ib. 10x Monomer Solution</u>**

1. Add 5.7 ml ddH2O into a 15 ml falcon tube. Let water cool down on the ice.
2. Prepare and mark 11x2ml empty tubes and store them on ice already.
3. Defreeze 38% Sodium Acrylate stock.

| Volume [ml] | Reagents | [Stock] | [Final] |
|---|---|---|---|
| 5.7 | ddH2O | | |
| 2.616 | N,N-dimethylacrylamide (DMAA) | >99% (102 g/100 ml) | 2.67 g/100 ml |
| 1.684 | Sodium Acrylate | 38% | 0.64 |
| 10 | **Final Volume** | | |

4. Vortex for 3 sec.
5. Put in each aliquot 900 µl of the Monomer Solution, store at -20°C. (Let polymerize the remaining rest of 100 µl in the falcon before discarding it.)
6. Refreeze unused Sodium acrylate. (Mark the cap with a red line for each defreezing procedure)

**II. Accelerator (TEMED solution)**

TEMED (10%): 50 µl TEMED (100%) + 450 µl ddH2O, store @ 4 °C

**IMPORTANT:** Get new TEMED when it turns yellow and stops stinking

**III. Digestion Buffer**

1. 8 M Guanidine HCl:
   - 19.106 g guanidine HCl (M=95.53 g/mol) in 25 ml ddH2O, store @ RT
2. 0.5 M EDTA (M=372.24g/mol)
   - 18.612 g EDTA in 65 ml ddH2O
   - Solubility of EDTA in water is 100 g/ml -> Already add about 1 g of NaOH pellet to increase solubility
   - Stir EDTA for solving.
   - While stirring adjust pH on 8 with NaOH 10 M. (About 2.5 ml in total, start with 0.5 ml)
   - After 2 h of stirring the EDTA solution should be clear.
   - Fill up the volume with dd$H_2O$ to 100 ml
   - filter sterile (filter size between 0.1 and 0.2 µm), store @ RT

3. 1 M Tris pH 8.0:
    - 12.114 g TRIS ultrapure (M=121.41 g/mol) in 70 ml ddH2O.
    - adjust pH with concentrated HCl to pH 8.0. (If you're using 1 M HCl, solve Tris in less than 70 ml of dd$H_2O$ water, as you need more than 30 ml of HCl to adjust to pH 8.0.)
    - Fill up the volume with dd$H_2O$ to 100 ml.
    - filter sterile (filter size between 0.1 and 0.2 µm) store @ RT
4. 10 M NaOH: **ATTENTION! Strong exothermic reaction!**
    - 4 g NaOH (MW: 40 g/mol) in 10 ml ddH2 on ice
5. 10 % Triton X-100: Put 10ml of Triton in a beaker and fill up with dd$H_2O$ to 100ml.
6. Digestion buffer stock solution:

| **Volume [ml]** | **Reagents** | **[Stock]** | **[Final]** |
|---|---|---|---|
| 35 | dd$H_2O$/PBS | | |
| 2.5 | Tris pH 8.0 | 1 M | 50 mM |
| 0.1 | EDTA pH 8.0 | 0.5 M | 1 mM |
| 2.5 | Triton X-100 | 10 % | 0.5 % |
| 5 | Guanidine HCl | 8 M | 0.8 M |
| 50 | **Final Volume** | | |

- adjust pH 8
- fill up to 50 ml.
- Store 9.9 ml aliquots @ -20 °C (amount: 5)

**Supplementary Protocol B: 10x Expansion of Platelets**

…………………………………………… Day 1…………………………………………….

**Preparation of coverslips**

1. Clean coverslips as mentioned in the methods part (day before day 1).
2. Decide how many samples you like to prepare and what size of coverslips you want to use. We use 24 mm for the unexpanded control samples and 12 mm for the 4x and 10x expanded samples
3. Cover flat backside of a 6- or 12- well plate with parafilm and place on it cleaned coverslips dried with $N_2$. (Morning Day 1)

**Remark: For 12mm coverslips we used a general volume of 100 µl, for 24 mm either 200 µl or 300 µl of the named reagents to immerse the whole surface.**

**<u>Step 1: Platelet preparation</u>**

**Step 1.1: Platelet preparation**

Prepare washed murine platelets (free of red blood cells) according to the standard procedure (see above) and adjust the platelet concentration to 400,000 platelets/µl Tyrode's buffer without calcium. Allow the platelets to rest at 37°C for 30 min.

**Step 1.2: Coating of coverslips**

1. Prepare 15 ml of 2 M Glycine in 1x PBS for coating the coverslips. You can store it at -20°C and just thaw it when required, for your next experiments.
2. Coat coverslips with 2 M Glycine for 10 min at RT.
3. Afterwards, remove Glycine and wash once with 1x PBS and with Tyrode's buffer with (activated platelets) or without $Ca^{2+}$ (for resting platelets), respectively.

**Step 1.3: Platelet activation**

4. **Choose the final concentration of platelets you want to have on the coverslips**. We used 50,000 platelets/µl for activated and 100,000 platelets/µl for resting platelets.
5. Prepare new tubes. You will now have to split up the platelet suspension into two volumes. To one of them you will later add thrombin for activation.
   Calculate the volume you have to take out of the high concentrated platelet suspension (400,000 µl) of tubes of point 25 and put it into the new tubes. Fill up with Tyrode's buffer

- $Ca^{2+}$ for resting and Tyrode's buffer + $Ca^{2+}$ to achieve the chosen concentration. NOTE: In point 6 you will add an additional volume of diluted Thrombin, so first calculate how much additional Thrombin volume you will add and then adjust the volume of the Tyrode's buffer + $Ca^{2+}$ for the activated platelets accordingly.

6. For the platelet activation:
   a. Dilute Thrombin from the stock (stock c=20 U/ml, kept at 4°C) to an intermediate stock with c = 0.1U/ml.
   b. Add Thrombin from the intermediate stock to the tube at a ratio of 1:10 with the platelet suspension you want to activate to have a final concentration of Thrombin of 0.01U/ml.
7. Pipette platelet solution carefully on the coverslip with round movements.
8. Let settle platelets for 30 min at 37°C in the incubator (or on the 37°C heating pad if you do not have an incubator).

**<u>Step 2: Fixation and Blocking</u>**

1. Prepare Glyoxal buffer by mixing the reagents in the table in a 15 ml falcon tube in the sequence as mentioned in the following table. Measure the pH-value of the solution. It should be around 5. If not, adjust with 2 M NaOH (about 160 µl might be required). Put in 37°C bath.

| **Volume [ml]** | **Reagents** | **[Stock]** | **[Final]** |
|---|---|---|---|
| 2.835 | dd$H_2O$ | | |
| 0.789 | Ethanol (100% pure) | 100% | 19.89% (wt/wt) |
| 0.313 | Glyoxal | 40% (wt/wt) | 3.15 % (wt/wt) |
| 0.03 | Acetic Acid | 99.5% | 0.7% (wt/wt) |
| 3.967 | **Final Volume** | | |

2. Remove the rest of the platelet suspension from the coverslips and add Glyoxal buffer. Incubate for 20 min at RT. (When you were using a heating pad you can also incubate for 2 min on heating pad (37°C) and 18 minutes at RT.)
3. Remove the fixation solution and wash 5 times with 1x PBS for 30 s, 1 min, 5 min, 10 min, 15 min.
4. Block slides with 5% BSA in 1x PBS for 1.5 h at RT.

**<u>Step 3: Labeling</u>**

**Step 3.1: Immunolabeling**

1. Prepare 1.5ml tubes with 5% BSA and add the fluorescently labeled primary antibody, so that your final primary antibody concentration in the BSA is 10 µg/ml (1:100). Work fast and cover the dyes and the tubes to prevent bleaching.
2. Give antibody solution on coverslips and incubate for 30min at 37°C in the incubator (or on the heating pad). **From now on, protect your sample from light.**
3. After incubation, wash 5 times with 1x PBS for 30 s, 1 min, 5 min, 10 min, 15 min.

**Step 3.2 Preparation of unexpanded samples**

4. Proceed with samples that you will expand directly with Step 4. For the unexpanded control samples follow the further steps for embedding.
5. Use Fluoroshield or Mowiol as embedding medium.
6. Put one drop of the embedding medium on a microscopic slide.
7. Flip the unexpanded coverslips with the platelets facing down directly on the medium.
8. Fix it with two drops of nailpolish. Let dry for 10-15 min and then seal it finally with nailpolish all around the coverslip.

**<u>Step 4: Linking</u>**

1. Solve the dry Acryloyl-X reagent in DMSO with a stock concentration of 10 mg/ml and aliquot 15 µl in 1.5-ml tubes. Store at -20°C.
2. Thaw Acryloyl-X aliquot and dilute stock (c = 10 mg/ml) to 0.1 mg/ml with 1x PBS by adding 1485 µl of 1x PBS to the 15 µl aliquot.
3. Place 100 µl Acryloyl-X solution on each coverslip with the samples you like to expand.
4. Incubate for 12 h (at least 6 h) overnight at RT in humidified chamber.

…………………………………………… Day 2……………………………………………

#### Step 5: Gelation

##### Step 5.1: Preparation of the bubbling process

1. The settings for the bubbling process will be installed under a hood.
2. Extend the tube of a $N_2$ gas tap to reach the hood and stick a pipette tip on the end. Take a 2ml tube filled with water to test the gas stream out of the tube. Reduce the stream to a continuous bubbling of 4-5 bubbles per second.
3. Take a 50 ml beaker and completely fill it with ice and water to prevent any free air spaces.
4. Install the beaker in the hood.
5. Install the test tube in the beaker so that it's covered around with water and ice. Arrange the tube so that the tip can insert the tube and switch on the gas. The bubbling should be consistent and not spilling out drops of the monomer solution. (By using the test tube for optimizing the setting you can save time later by just replacing the test tube with the tube containing the monomer solution.

##### Step 5.2: Gelation

1. Get a box with ice.
2. Cover the flat backside of a 6- or 12-well-plate with parafilm and put it on the patted ice (flat surface).
3. Prepare 10x Monomer solution: Bubble at RT for 40 min with N2 ** **(See Step 5.1)**
4. KPS-Solution (Potassium Persulfate) has to be always prepared freshly. Dilute 0.18 g KPS in 5 ml $ddH_2O$ and put it on ice.
5. After 40 mins of bubbling, add 100 µl KPS to Monomer solution and continue bubbling with $N_2$ for 15 min.
6. Take your coverslips covered with the Acryloyl-X solution from the day before. Take out only 95 ul solution or the platelets might dry. Wash them 2x times with PBS, each interval of washing spaced 5 mins apart.
7. Discard the PBS, applied in the washing step, let dry but not completely.
8. Add 4 µl TEMED (100%) to the KPS-Monomer-solution and vortex briefly. From now on, act quickly.

| Volume [µl] | Reagents | [Stock] | [Final] |
|---|---|---|---|
| 900 | Monomer solution | | |
| 100 | KPS (initiator) | 0.036 g/ml | 0.0036 g/ml |
| 4 | TEMED (Accelerator) | 100% | 0.4% |
| 1004 | **Final Volume** | | |

9. Pipette 50 µl of the solution onto parafilm.
10. Cover the drop of gelation solution with the coverslip – cells facing down - and make sure the gelation solution is evenly distributed beneath the glass
11. Take the $N_2$ pumping box and put the coverslips containing plate into the box with the lid partially on. Now put the $N_2$ nozzle into the box with the lids closed and pump $N_2$ for about 2-3 minutes. Then seal the box(es).
12. Protect sample from light and let gelatinize for 48 h in a humid chamber at 4°C.
13. Gelation is finished, when coverslip with the gel can be easily removed from parafilm.

……………………………………………… Day 4………………………………………………

**Step 6: Digestion**

1. Thaw digestion buffer and preheat oven (50°C).
2. Digestion Buffer with Proteinase K:

| Volume [ml] | Reagents | [Stock] | [Final] |
|---|---|---|---|
| 9.9 | Digestion Buffer | 1x | 1x |
| 0.1 | Proteinase K | 600 *units/ml | 8 U/ml |
| 10 | **Final Volume** | | |

3. Transfer gel into small petri dish.
4. Add to each petri dish Digestion Buffer with Proteinase K so that Gel is covered well. 1.5 ml at least, more is possible.
5. Put dishes in oven for at least 12 h @ 50°C in humid chamber (closed)

……………………………………………… Day 5………………………………………………

**Step 7: Expansion**

1. For X10 Gel Expansion you should use dishes with 12 cm diameter. Put the gel into those bigger dishes. If the gels sticks in the small petridish, add some water and move in circles.
2. Wash gels with excess volume of dd$H_2O$ 3-5 times a day for 0.5 h- 2 h. Last washing step should be overnight, so that the gel can fully expand.
3. Remove the water on the next morning and measure and note diameter for the macroscopic expansion factor.

……………………………………………… Day 6………………………………………………

**Step 8: Imaging**

Immobilization of gel by Poly-D-Lysine coating of coverslip

1. Dry cleaned glass coverslips (see methods) with $N_2$ and place them on the flat backside of a 6-well plate covered with parafilm.
2. Dilute thawed Poly-D-Lysine to 0.25 mg/ml by adding 1.5ml dd$H_2O$ to the stock solution.
3. Put 500 µl on each coverslip and incubate at 37°C for 1h.
4. Wash twice: first time add 300 µl dd$H_2O$ without removing the solution. Incubate for 3 min. Then remove the solution completely and add 300 µl dd$H_2O$.
5. Store the coverslips covered with this dd$H_2O$ for up to 14 days in the fridge. Before using the coverslips dry them with nitrogen and check if they are still clean.
6. Mount dried coverslip in an imaging chamber (eg. Attofluor cell chamber)
7. Cut a piece of gel, make sure you know the gel orientation. The side with the platelets should face towards the coverslip.
8. Balance the piece of gel on a spatula and remove excess water with a tissue.
9. Place the gel carefully on the dry Poly-D-Lysine coated coverslip, let it rest for 5 minutes.
10. Add water to the imaging chamber.
    **If the gel is not attached to the surface, it will start swimming away. You can dry the gel again and try to attach it to a fresh, dry Poly-D-Lysine coated coverslip.**
11. Now you can start imaging!

**Supplementary Protocol C: 4x Expansion of Platelets**

……………………………………………… Day 1………………………………………………

The first day (from platelet washing, spreading, fixing, blocking, immunolabelling and then linking) stays the same as described in **Supplementary Protocol B**.

……………………………………………… Day 2………………………………………………

**Step 1: Gelation**

1. Get a box with ice and cover the flat backside of a petri plate with parafilm. Put everything on ice, this provides more time for handling the gels.
2. Wash the coverslips covered with Acryoly-X from the day before twice with PBS 1x for each 5min.
3. For 10% APS: Either dilute the frozen stock (stored at -20°C for 1 month) from 50% to 10% (10 µl APS 50% plus 40 µl dd$H_2O$) or prepare a new stock. (5 g APS and fill up with dd$H_2O$ to 10ml)
4. Prepare TEMED: Either use 10% stock or dilute 100% stock to 10% (10 ml 100% TEMED in 90 ml dd$H_2O$, store in dark and at 4°C).
5. Prepare gelation solution on ice. First add water, then APS, then TEMED to monomer solution. Vortex 3 sec. Keep the monomer solution on ice.

| **Volume [µl]** | **Reagents** | **[Stock]** | **[Final]** |
|---|---|---|---|
| 20 | APS (initiator) | 10% | 0.2% |
| 20 | TEMED (Accelerator) | 10% | 0.2% |
| 950 | Monomer solution | | 98.6% |
| 10 | ddH2O | | |
| 1000 | **Final Volume** | | |

6. Immediately pipette 50 µl of gelation solution onto parafilm.
7. Now immediately cover the drop of the gelation solution with the coverslip – cells facing down.
8. Make sure the gelation solution is evenly distributed beneath the glass.
9. Move lid of the well plate covered with the upper part in a humid box.
10. Protect sample from light and move to 37° C incubator for 1.5 h for gelation.
11. Gelation is finished when the gel can be easily removed from the parafilm.

……………………………………………… Day 3………………………………………………

**Step 2: Digestion**

Digestion is performed as described in **Supplementary Protocol B - Step 6** for at least 8 h or overnight at RT in a humidified chamber.

……………………………………………… Day 4………………………………………………

**Step 3: Expansion**

1. Transfer gels into big petri dish (6 cm diameter) and cover with an excess of dd$H_2O$.
2. Exchange dd$H_2O$ every 0.5 h till 2 h 5 times. Last step over night.

### Major Resources Table

#### Animals (in vivo studies)

| Species | Vendor or Source | Background Strain | Sex | Persistent ID / URL |
|---|---|---|---|---|
| C57Bl/6J mice | Charles River | C57Bl/6J | M/F | Jax number: 000664 |

#### Antibodies

| Target antigen | Vendor or Source | Catalog # | Working concentration | Lot # (preferred but not required) | Persistent ID / URL |
|---|---|---|---|---|---|
| IgG Anti-GPIIb/IIIa Antibody (MWReg30) | In-house produced | N.a. | 10 µg/ml | N.a. | N.a. |
| IgG Anti-GPIIb/IIIa Antibody (JON6) | In-house produced | N.a. | 10 µg/ml | N.a. | N.a. |
| IgG Anti-GPIX Antibody p0p6 | In-house produced | N.a. | 10 µg/ml | N.a. | N.a. |

#### Data & Code Availability

| Description | Source / Repository | Persistent ID / URL |
|---|---|---|
| A selection of typical images, the code for simulation of platelet receptor distributions and a video protocol of 10x ExM procedure | ZENODO | doi: 10.5281/zenodo.4117555 |

#### Other

| Description | Source / Repository | Persistent ID / URL |
|---|---|---|
| Round coverslips Ø12 mm, #1.5 | Menzel | https://www.fishersci.de/shop/products/11846933/11846933# |
| Round coverslips Ø24 mm, #1.5 | Plano GmbH | N.a. |
| Chloroform | Sigma Aldrich (472476) | https://www.sigmaaldrich.com/catalog/product/sigald/472476 |
| Sodium-hydroxide | Roth (6771) | https://www.carlroth.com/at/en/reagents-for-the-determination-of-nitrogen-acc-to-kjeldahl/sodium-hydroxide/p/6771.1 |
| Ethanol | Sigma Aldrich (34852) | https://www.sigmaaldrich.com/catalog/product/sigald/34852 |
| Glycine | AppliChem (A1067) | https://www.applichem.com/en/shop/product-detail/as/glycin-fuer-die-molekularbiologie/ |
| Glyoxal | Sigma Aldrich (128465) | https://www.sigmaaldrich.com/catalog/product/sial/128465 |
| Acetic acid | Roth (7332.1) | https://www.carlroth.com/de/en/lewis-acids/acetic-acid/p/7332.1 |
| Bovine serum albumin (BSA) | Sigma (A3983) | https://www.sigmaaldrich.com/catalog/product/sigma/a3983 |
| Alexa Fluor 488 | Thermo Fisher (A20181) | https://www.thermofisher.com/order/catalog/product/A20181 |
| Alexa Fluor 594 | Thermo Fisher (A20185) | https://www.thermofisher.com/order/catalog/product/A20185 |
| Acryloyl-X | Thermo Fisher (A20770) | https://www.thermofisher.com/order/catalog/product/A20770 |
| Acrylamide (AA) | Sigma Aldrich (A9099) | https://www.sigmaaldrich.com/catalog/product/sigma/a9099 |
| N,N′-methylenebisacrylamide | Sigma Aldrich (M7279) | https://www.sigmaaldrich.com/catalog/product/sigma/m7279 |
| Sodium acrylate (SA) | Sigma Aldrich (408220) | https://www.sigmaaldrich.com/catalog/product/aldrich/408220 |

| | | |
|---|---|---|
| Ammonium persulfate (APS) | Roth (9592.3) | https://www.carlroth.com/de/en/ammonium-salts-nh4/ammonium-peroxydisulphate/p/9592.1 |
| Tetramethylethylenediamine (TEMED) | Roth (2367.3) | https://www.carlroth.com/de/en/page-reagents/temed/p/2367.3 |
| Proteinase K | Sigma Aldrich (P4850) | https://www.sigmaaldrich.com/catalog/product/sigma/p4850 |
| EDTA | Sigma Aldrich (ED2P) | https://www.sigmaaldrich.com/catalog/product/sial/ed2p |
| Triton X-100 | Thermo Fisher (28314) | https://www.thermofisher.com/order/catalog/product/28314#/28314 |
| Guanidine HCl | Sigma Aldrich (50933) | https://www.sigmaaldrich.com/catalog/product/sigma/50933 |
| Poly-D-lysine (02102694) | MP Biomedicals (02102694) | https://www.mpbio.com/eu/poly-d-lysine-hydrobromide-mw-4-000-15-000 |
| N,N-dimethylacrylamide (DMAA) | Sigma Aldrich (274135) | https://www.sigmaaldrich.com/catalog/product/aldrich/274135 |
| Potassium persulfate (KPS) | Sigma Aldrich (206224) | https://www.sigmaaldrich.com/catalog/product/SIGALD/216224 |
| 4-colour fluorescent microspheres (TetraSpeck) | Thermo Fisher (T7279) | https://www.thermofisher.com/order/catalog/product/T7279 #/T7279 |
| Apyrase | Sigma Aldrich (A6410) | https://www.sigmaaldrich.com/catalog/product/sigma/a6410 |
| PGI2 | Sigma-Aldrich (P6188) | https://www.sigmaaldrich.com/catalog/product/sigma/p6188 |
| Thrombin (20U / 22.23 mg lyo) | Roche Diagnostics (10602400001) | https://www.sigmaaldrich.com/DE/de/product/roche/10602400001 |
